# TRPV4 Dominates High Shear-Induced Initial Traction Response and Long-Term Relaxation Over Piezo1

**DOI:** 10.1101/2025.02.10.637570

**Authors:** Mohanish K. Chandurkar, Manli Yang, Majid Rostami, Sangyoon J. Han

## Abstract

Modulation of endothelial traction is critical for the responses of endothelial cells to fluid shear stress (FSS), which has profound implications for vascular health and atherosclerosis. Previously, we demonstrated that under high FSS, endothelial cells rapidly increase traction forces, followed by relaxation, with traction aligning in the flow direction. In contrast, low shear preconditioning induces a modest short-term increase in traction (<30 min), followed by a secondary long-term (>14 hr) rise, with traction/cells aligning perpendicular to the flow. The upstream mechanosensors driving these responses, however, remain unknown. Here, we sought the roles of Piezo1 and TRPV4 ion channels in shear-induced traction modulation. We report that HUVECs with Piezo1 silencing reduced the initial traction rise in half under high FSS compared to those by WT cells, while not affecting the traction modulation in response to low FSS or traction/cell alignment to the flow direction. Conversely, cells with siTRPV4 fully abrogated the initial traction rise, as well as alignment of traction and cells, in response to both high and low FSS conditions. Dual inhibition of Piezo1 and TRPV4 further impaired both initial and long-term traction under high FSS. Interestingly, dual-inhibited cells displayed larger initial traction responses to low FSS compared to control cells, suggesting the involvement of alternative calcium-independent pathways that become dominant when both ion channels are nonfunctional. Additionally, either ion channel inhibition led to secondary long-term traction increase even under high FSS condition. These findings suggest that while both Piezo1 and TRPV4 channels contribute to shear mechanotransduction, TRPV4 plays more dominant role than Piezo1 in mediating the initial traction rise and sustaining long-term relaxation under high or low shear stress, highlighting their critical and distinct contributions to endothelial mechanotransduction and remodeling.

## Introduction

Mechanotransduction in endothelial cells (ECs) in response to fluid shear stress (FSS) is critical in maintaining barrier functions, vascular integrity, and for regulating inflammation and disease progression^1,2^. FSS initiates a range of signaling and transcriptional pathways, from immediate responses to long-term adaptations^3^. Depending on the magnitude and pattern of FSS, ECs can adopt either protective or pathological phenotypes. High, steady FSS aligns with atheroprotective responses, promoting cell alignment in the direction of flow and maintaining vascular integrity^4,5^, while low or oscillatory FSS is often associated with atheroprone environments, leading to perpendicular cell alignment, increased inflammation, and compromised endothelial function. Understanding how ECs sense and respond to these mechanical cues is essential for uncovering the mechanistic basis of endothelial adaptation and maladaptation under varying shear conditions.

Piezo1, a mechanosensitive calcium ion channel, plays a fundamental role in the development and maintenance of the cardiovascular system and has been implicated in vascular biology and pathology, including atherosclerosis^3,6–9^. Located on the apical surface of endothelial cells (ECs), Piezo1 is activated by mechanical stimuli such as fluid shear stress (FSS), triggering calcium (Ca²⁺) influx^10^. This Ca²⁺ influx is associated with integrin activation^3,11,12^ and endothelial cell alignment in response to flow^13,14^. Accordingly, Piezo1 is critically involved in flow-induced production of atheroprotective signaling protein KLF2/4^15^ and nitric oxide (NO), which are essential for blood pressure regulation and vascular tone^16^.

Similarly, transient receptor potential cation channel V4 (TRPV4), belonging to a transient receptor potential (TRP) family ^17^, is also linked with regulation of blood flow and by shear shear-induced activation to control vasodilation and remodeling^18,19^. As in Piezo1, once activated by mechanical cues such as shear stress^17^, stretch^20^, temperature changes, and osmotic pressure, TRPV4 facilitates Ca²⁺ influx that initiates downstream signaling pathways including NO synthesis^21^, cytoskeletal dynamics, focal adhesion (FA) remodeling, and ultimately, traction force (TF) generation and alignment behaviors in ECs^22^. However, differential regulation of endothelial mechanotransduction from TRPV4 compared to Piezo1 has been unclear.

TFs serve as a physical readout of endothelial mechanotransduction, providing direct insights into how cells sense and respond to shear stress. These forces arise from the dynamic interactions between the cytoskeleton, FAs, and the extracellular matrix, enabling ECs to modulate their shape, alignment, and function in response to mechanical stimuli. Our previous findings demonstrated that under high FSS, endothelial cells exhibit a rapid increase in traction forces followed by relaxation, with traction aligning in the direction of flow^23^. In contrast, low shear preconditioning induces a modest initial increase in traction, followed by a secondary long-term rise, with traction and cells aligning perpendicular to the flow^23^. This differential traction response suggests that distinct mechanosensitive pathways regulate endothelial adaptation to varying shear conditions. However, the upstream regulators responsible for modulating these forces remain unclear.

Given the established roles of Piezo1 and TRPV4 in shear-induced calcium signaling, we sought to investigate their contributions to endothelial TF modulation and cell alignment under different shear stress conditions. Using siRNA to selectively inhibit Piezo1 and TRPV4 in human umbilical vein endothelial cells (HUVECs), we quantify TF generation and alignment dynamics under direct high FSS and low FSS preconditioning. Our results reveal that Piezo1 primarily regulates traction responses and cell alignment under high FSS, while TRPV4 controls the rapid initial traction response and alignment under both high and low FSS. Dual inhibition experiments suggest that these channels together contribute to the magnitude and direction of traction forces, with their inhibition leading to distinct alterations in traction and alignment based on the level of FSS applied. These findings provide new insights into the role of mechanosensitive ion channels as primary FSS sensors in endothelial cells and offer a mechanistic basis for understanding how ECs differentiate between atheroprotective and atheroprone shear environments.

## Material and Method

### Construction of a Flow System with Traction Microscopy

A rectangular flow chamber was used as previously fabricated^23^. Briefly, the chamber has a dimension of 38ξ5ξ0.8mm for the chamber’s inner volume and was manufactured using polycarbonate via CNC machining. This flow chamber was attached to the bottom coverslip via adhesive tape (3M). Before attachment to the flow chamber top, the cover glass was prepared to contain a silicone gel with fluorescent markers for traction measurement (Fig. 1 A). Coating the cover glass with a silicone gel and coating the gel with fluorescent beads were done as previously done ^24^. The assembled flow chamber was integrated into the flow system that consists of a peristaltic pump (model no. NE-9000; New Era Pump Systems, Inc.), a pulse dampener, and a reservoir (Fig. 1D). The pulse dampener was used to eliminate the pulsation in the flow due to the pump and to ensure a continuous flow of cell media without any air bubbles. A glass bottle was used as a reservoir, which has three holes for the inlet, outlet, and CO_2_. The CO_2_ concentration and the temperature inside the chamber were maintained by enclosing the media inside the CO_2_ incubator. The pump was operated using LabVIEW for flow profile programming flexibility. The flow system was maintained at 37 °C and 5% CO_2_ throughout the experiments.

**Figure 1.**
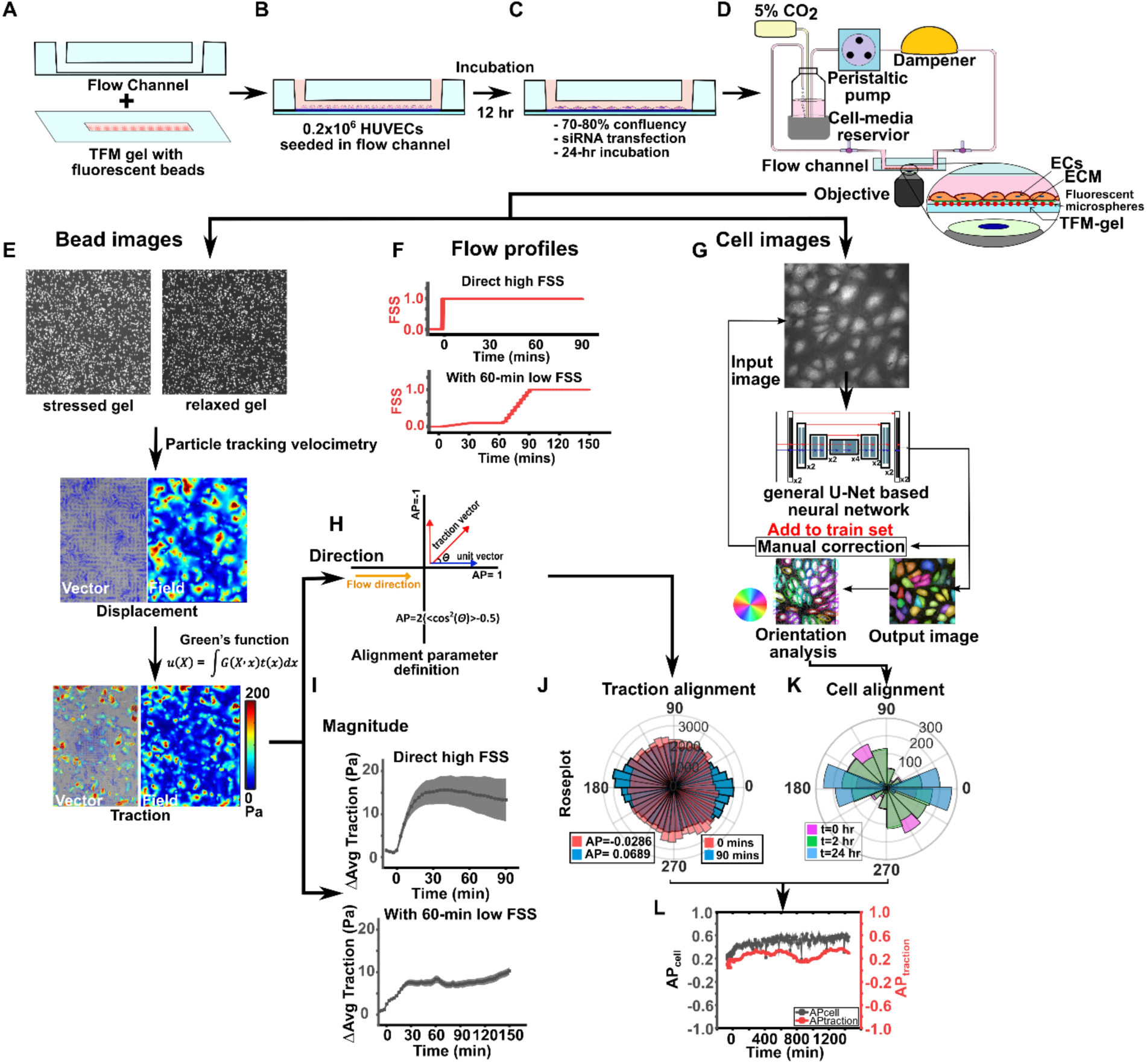
Flow system setup and traction microscopy (TM) workflow and analysis. (A) Flow channel and glass coverslip with TM gel and fluorescent beads for TM imaging. (B) HUVECs seeded in the flow channel coated with fibronectin and incubation for 12 hours to reach 70-80% confluency. (C) Transfection of cells using siRNA for piezo1, TRPV4 and both with further incubation for 24 hours. (D) Illustration of the flow system with peristaltic pump, pulse dampener, media bottle maintained at 5% CO_2_ and 37 °C, the flow channel setup on microscope for live TM imaging and 20x objective. (E) Illustration of the TM gel with stressed and relaxed configurations along with representative bead images on the right. The bead tracking using particle tracking velocimetry between stressed and relaxed gel followed by measurement of displacement field and vectors. Traction field and vectors calculated using green’s function for known thickness of 24 µm and stiffness of the gel 2.6 kPa. (F) Flow profiles used for the FSS application (up) Direct high FSS, (down) With 60-min low FSS. (G) Cell orientation analysis framework consisting of Cellpose 2.0 neural network that uses a standard U-net backbone, which down sample and up sample the feature maps along with skip connection between layers, with a human-in-loop procedure for manual training and correction. Output image after segmentation prediction was analyzed for orientation in custom developed python code for alignment parameter with angular distribution of the segmentation. (I) Change in traction magnitude plotted over a timescale for the applied FSS of Direct high and 60-min low FSS. (J) Polar-histogram distribution of force vectors at timepoints 0 mins (pink) and 90 mins (blue). (K) Polar-histogram distribution of cell orientation at timepoints 0 hours (pink), 2 hours (green), and 24 hours (blue). (L) Representation of time-series plot of alignment parameter, AP_cell_ (gray) and AP_traction_ (red) as a function of time from each representative time-lapse image recorded for FSS.

### Cell Culture, Silencing, and Seeding to the Flow Chamber

The HUVEC cell line was purchased from Lonza (Cat. No. C2519A). The cells were used from passages 3–7 for the experiments and were cultured in endothelial cell growth medium-2 (EGM-2) Bulletkit (Lonza Bioscience) on a pretreated tissue culture six-well plate (VWR international). For the flow experiments, 0.2×10^6^ cells/mL were seeded on the soft gel-coated coverslip and stored in a CO_2_ incubator for 12 hours to reach 70-80% confluency (Fig. 1 B). The HUVECs were then transfected with siRNAs of Piezo1 (ON-TARGET plus Human Piezo1 siRNA, Catalog number: J-020870-09-0005, Dharmacon), and TRPV4 (ON-TARGET plus Human TRPV4 siRNA, Catalog number: J-004195-08-0005, Dharmacon), as per manufacturers protocol using lipofectamine RNAiMAX transfection reagent. The cells were allowed to form confluency and are incubated for at least 24 hours to achieve transfection efficiency as per manufacturers’ recommendation (Fig. 1C). The cells were treated with CellTracker^TM^ Fluorescent dye (Catalog number: C7025, ThermoFisher scientific) 1 hour before the start of the experiment to observe cell confluency, orientation, and alignment. Functionalization of the gel with fibronectin for cell adhesion was performed as previously done^25^.

### mRNA Measurement Using Real-Time PCR

qRT-PCR was performed to validate the knockdown results of the piezo1 and trpv4 genes in HUVECs. 2×10^5^ HUVECs were seeded on each well of a 6-well plate. After 24 hours, cells were incubated with or without siRNA, ON-TARGETplus Non-targeting Control Pool (Catalog No. D-001810-10-05), ON-TARGETplus Human PIEZO1 siRNA, ON-TARGETplus Human TRPV4 siRNA. After 24 hours, the medium was replaced with fresh culture medium. Another 24 hours later, RNA was isolated using the Qiagen Micro kit (Catalog No. 74004). Then, between 200 and 500 ng of RNA from each sample was used to perform reverse transcription using the iScript cDNA Synthesis Kit (Catalog No.1708890) to obtain cDNA. Subsequently, qPCR was performed using 20 ng of cDNA with the SYBR Green method.

### Shear Flow Experiments

For the shear flow experiments, the calculations for FSS were performed using the Poiseuille law modified for parallel-plate flow channels^26^:

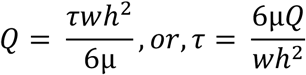

where Q is the desired flow rate, τ is the wall shear stress on the gel and cell surface, μ is the viscosity of the cell media (0.72 mPa·s), w is the width of the flow channel (5 mm), and h is the height of the channel (0.8 mm). Accordingly, to apply 1-Pa FSS, for example, the flow rate of 44.4 mL/min was designed and applied. We have two different flow profiles: 1) direct high FSS, and 2) with 60-min low FSS (Fig. 1 F). For direct high FSS, the FSS of 1 Pa was directly applied with the onset of the flow from the static condition. Our selection of 1 Pa as the high FSS level was informed by the literature^27–29^ indicating that shear stress within the 1- to 2-Pa range is sufficient to induce atheroprotective gene expression and anti-inflammatory responses in vitro, paralleling physiological conditions in larger vessels. This level was also chosen for its practical compatibility with our experimental setup, balancing the need for physiological relevance with technical feasibility.

For the flow with 60-min low FSS, the FSS started from 0 to 0.01 Pa and was ramped every minute with 0.0031 Pa to reach 0.1 Pa at t = 30 min. It was followed by a constant 0.1-Pa FSS for 30 min until t= 60 min, after which the FSS was further increased from 0.1 to 1 Pa with a linear ramp until t= 90 min. For flow profiles, after reaching 1 Pa, the flow was maintained at 1 Pa until t= 24 h. For short-term imaging, which is 90, and 150 min, for direct high FSS, and 60-min low FSS, respectively, the time interval of 3 min was used. For a long-term imaging until 24 h, the time interval of 10 min was used. Justification for this time frame determination and the flow control using a programmable pump are described in^30^.

### Live Cell Imaging

The beads in the flow chamber were imaged via a spinning-disk confocal microscope (Yokogawa CSU-X1, built on Nikon Ti-S) with 20 objective and 642-nm laser excitation before and after the flow application at every 180s (Fig. 1 E). At the same time, differential interference contrast (DIC) images and fluorescent cell images were taken to observe the cell shapes, confluency, and orientation (Fig. 1 G). The reflected light from SD Confocal is acquired by the ORCA-Flash sCMOS camera. Image acquisition was controlled by Metamorph software. During imaging, the flow chamber was kept inside a stage-top incubator (OKO Laboratory) using a sealing putty to ensure stable imaging without drifting. For TM imaging, after imaging in cell presence, HUVECs were removed from the substrate surface by flowing 0.25% trypsin into the chamber, and then the bead positions in relaxed gel configuration were imaged via far-red laser again (Fig. 1 E).

### Traction Reconstruction and Analysis

The acquired images of the beads on the deformed gel (when HUVECs were present on the substrate) and of the reference configuration of the relaxed gel (after removing HUVECs) were processed for traction reconstruction using traction microscopy (TM) as described in our previous publications (Fig. 1 E, I)^31–34^. Detailed TM parameters are described in reference^30^. After traction reconstruction, the magnitude of the traction was plotted over timescale to monitor the change in the traction before and after FSS (Fig. 1I). For visualization of the direction of orientation of the traction vectors, the generated force field was used to calculate the angular distribution of the force alignment using a custom-written code in MATLAB. The vector alignment to the flow direction was calculated using the alignment parameter, AP_traction_ = 2(<cos^2^θtraction>-0.5), where θ_traction_ represents the angle of a traction vector to the flow direction, which is an x-direction in our configuration (Fig. 1 H). The average of the distribution of angles for the force alignment was denoted using < >. For AP_traction_, only prominent traction vectors were used, by subsampling with Otsu thresholding, for the average calculation to avoid noise-like effect from small traction vectors.

### Cell Segmentation, Training, and Orientation Analysis

The Cellpose 2.0^35^ was used with a human-in-loop model to predict cell segmentation. The training was done on the bright-field images or fluorescent dye cell images of HUVECs for each subset of images and the prediction was further improved with manual training and correction. The models were trained from scratch for 100 epochs with a batch size of 15, with a weight decay of 0.0001 and a learning rate of 0.1. For all the segmentation predictions, images with less than five predicted cells were excluded for orientation calculation. The orientations of the segmented areas were determined by the major axes of ellipses fitted to each segmentation. The cell alignment to the flow direction was calculated using the alignment parameter, AP_cell_ = 2(<cos^2^θ_cell_>0.5), where θ_cell_ represents the angle of the segment’s major axis to the flow direction (Fig. 1 G). For visual inspection of the orientations of traction vectors or cells’ major axes, the polar histogram was plotted using the MATLAB function, “polarhistogram” (Fig. 1 K, L). The time series analysis of the AP for cell and traction were plotted over a timescale of 24 hours (Fig. 1 M).

### Statistical Analysis

The data were obtained from three replicate experiments and five different positions in each experiment, unless stated otherwise. Error bars in all figures represent the standard error of the mean and the analysis was done using OriginLab. To determine a significant difference between groups, we used one-way ANOVA with Tukey’s post hoc test. To determine if the analyzed data, for example, change in traction or change in AP, are significantly more than zero, we used one sample Student’s t test per group. ANOVA with Tukey’s post hoc test and Student’s t test were done in JMP-pro software.

The Granger causality (GC) test was performed to determine whether AP_traction_ functionally causes AP_cell_ or vice versa as discussed in ref ^30^. Briefly, we analyzed the causality in both directions, that is, AP_cell_ GCs AP_traction_ and AP_traction_ GCs AP_cell_, from which a significance of the causality, that is, P values, was produced. In each experiment condition, the P values were combined using the HMP to determine the global representative significance of each causal direction. A global representative significance of <0.05 was considered significant. Furthermore, we checked if the two sets of GC P values, representing the two GC directions, were significantly different. Since distributions of P values were far from being Gaussian, the usual two-sample t test could not be used. We instead used the Mann–Whitney test. These P values are further combined using HMP.

## Results

### Piezo1 modulates ECs traction response under direct high FSS but not under low FSS and involved in traction and cell alignment

Previously, using flow-TM experiments (as illustrated in Fig. 1), we have shown that, in wild-type (WT) HUVECs exposed to direct high FSS, traction exhibits a distinct temporal profile. In the short-term (Fig. 2A, *gray curve*), traction forces increase rapidly within the first few minutes of FSS application, reaching a peak within 5-10 minutes, followed by a steady phase over the next hour. This quick traction rise, and stabilization indicates an initial, robust mechanotransduction response to high FSS in WT ECs. In the long-term response (Fig. 2B, *gray curve*), traction, initially risen, rapidly decreases below the baseline, showing a relaxation phase that suggests adaptation to prolonged high FSS. When wild-type HUVECs undergo 60-minute low FSS preconditioning before high FSS application the traction increase is more gradual and modest compared to direct high FSS, with a slower rise and lower peak (Fig. 2C, *gray curve*), indicating that low shear preconditioning reduces the initial flow sensitivity of wild-type ECs. Over the long term, however, the traction force under this preconditioned regimen shows a huge secondary rise for over 10 hours before slowly declining (Fig. 2D, *gray curve*)^30^. These trends in wild-type cells provide a baseline for understanding the altered traction responses observed in Piezo1-inhibited cells.

**Figure 2.**
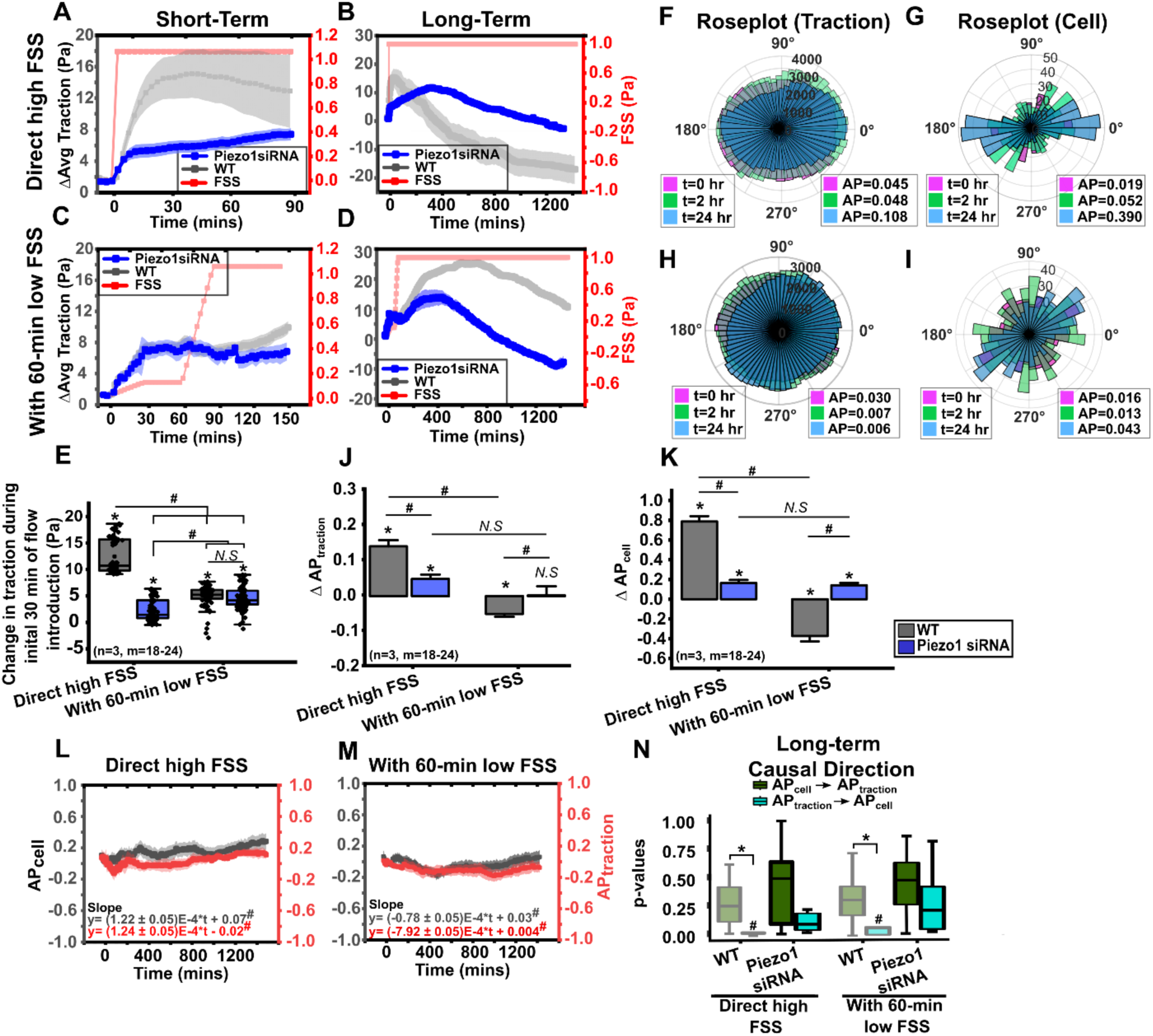
Piezo1 inhibition impairs early traction response to direct high FSS, prolonging long-term traction relaxation and also impairs alignment under high and low FSS. (A, B, C, D) Average traction of HUVECs silenced with Piezo1 siRNA (blue) for short-term (A, C) and long-term (B, D) under direct high FSS (A, B) and 60-min low FSS (C, D). Wildtype condition shown in (grey) and FSS in (red) (N=1, M=7-9). Error bar means ± SE. (E) A boxplot of early rise in traction within 30 min of FSS (N=3, M= 18-24). (F, H) Polar distribution of traction vectors at t=0 hr (magenta), t=2 hr (green) and t=24 hr (blue) for each condition. (G, I) Polar distribution of cell orientation at t=0 hr (magenta), t=2 hr (green) and t=24 hr (blue) for each condition. (J, K) Bar plot of changes in ΔAP_traction_ and ΔAP_cell_ for wild-type (*gray*) and Piezo1siRNA (*blue*) under direct high FSS and 60-min low FSS conditions. (N=3, M= 18-24). (L, M) Time-series plot of alignment parameter, AP_cell_ (gray) and AP_traction_ (red) as a function of time from each representative time-lapse images recorded for direct high FSS (L), and 60-min low FSS (M). #*P*<0.05 tested with t-test for the slope of the line, compared against null hypothesis that the true slope is zero. N represents the number of independent experiments and M represents the position observed. \**P*<0.05 tested with one-sample t test, that is, compared against zero. # *P*<0.05 tested with one-way ANOVA with Tukey’s post hoc test. (N) Boxplots of P values from Granger causality (GC) tests for long-term imaging for all conditions with causal direction that “AP_cell_ GCs AP_traction_” (green) and “AP_traction_ GCs AP_cell_” (blue). #P < 0.05 tested with HMP-combined GC P values, *P < 0.05 tested with Mann–Whitney U test between the two-direction GC test P values. Sample sizes for each condition are N = 4, 10, 5, 10, for direct high FSS (WT), direct high FSS (Piezo1 siRNA), 60-min low FSS (WT), 60-min low FSS (Piezo1 siRNA) respectively.

To determine the role of Piezo1 channel in regulation of flow-mediated traction (Fig. 1E), we measured the traction in HUVECs treated with Piezo1 siRNA under two conditions: direct high FSS and high FSS following 60 minutes of low FSS preconditioning (Fig. 1F, *top* and *bottom*, respectively). Knock-down efficiency of Piezo1 silencing (∼89%) was confirmed by qPCR (Fig. S1). Upon exposure to direct high FSS, Piezo1-inhibited cells displayed a noticeable initial increase in traction within the first 5 minutes (Fig. 2A, Fig. S3A), where the increase in traction was significantly lower than that of control cells (Fig. 2A, E). This initial rise was followed by a more gradual increase. In the long term, Piezo1-inhibited cells exposed to direct high FSS showed a slower, steady increase in traction over several hours, eventually reaching a maximum traction level that was lower than the global maximum observed in wild-type cells (Fig. 2B). This was followed by a relaxation phase also slower than that of the control ECs (Fig. 2B, S3B), which was similar to the secondary rise shown in WT cells under 60-min low FSS.

In contrast, in the condition with 60-minute low FSS preconditioning followed by high FSS, Piezo1-inhibited cells exhibited a similar trend to wild-type cells in the short-term response (Fig. 2C, E, S3C): *i.e.*, traction initially rose within the first few minutes and then plateaued, showing no further significant change even as FSS increased. This finding suggests that Piezo1-mediated traction is insensitive to low FSS in the short term. Over the long term, Piezo1-inhibited cells under this preconditioned regime showed a secondary increase in traction similar to wild-type cells, but the maximum traction achieved was still lower than in control conditions (Fig. 2D, S3D). Additionally, the relaxation phase in Piezo1-inhibited cells occurred more rapidly, dropping below baseline traction levels within 24 hours, indicating a faster return to baseline compared to wild-type cells.

Our recent evidence has shown that traction alignment functionally causes cell alignment^30^ To examine Piezo1’s role in the shear-induced traction alignment, we analyzed the angular distribution of traction vector fields (Fig. 2F, H, J). Before the onset of the FSS, the distribution of the traction vectors showed mostly isotropic distribution (Fig. 2F, H, *pink*). Under direct high FSS, the vector orientation showed a shift towards the flow direction as a function of time (Fig. 2F, *green* and *blue*) whereas under 60-min low FSS condition, the traction field showed a slight shift toward both the direction parallel and perpendicular to the flow (Fig. 2H, *green* and *blue*). Analysis of the change in the alignment parameter (AP_traction_, as described in Fig. 1K) of all experiments confirmed a positive shift in the alignment of Piezo-inhibited HUVECs under direct high FSS condition after 24 hours with the magnitude of the change smaller than that of control (Fig. 2J). This suggests Piezo1’s partial effect in high FSS-triggered traction. In response to 60-min low FSS condition, however, HUVECs showed an insignificant shift in AP_traction_, which was different from the negative change in AP_traction_ exhibited in WT cells under low FSS (Fig. 2J). Thus, Piezo1 inhibition impairs traction’s perpendicular alignment to the flow under low FSS while despite traction magnitude reduction.

The analysis of cell orientations and alignment parameters (AP_cell_) using deep neural network (Fig. 1G, I, K, S4) in Piezo1-inhibited HUVECs under different shear conditions reveals key differences in their response to direct high FSS and 60-min low FSS preconditioning. Under direct high FSS, the angular distribution of cell orientations (Fig. 2G) shows significant alignment in the flow direction, albeit less prominently compared to wild-type cells. Conversely, after 60-min low FSS preconditioning, no notable alignment was observed, with cell orientations remaining largely isotropic (Fig. 2I). Quantitative analysis of alignment parameters (ΔAP, Fig. 1L) further underscores these differences (Fig. 2K). Under direct high FSS, Piezo1-inhibited cells exhibit a significantly attenuated increase in both ΔAP_cell_ and ΔAP_traction_ compared to those exhibited by WT cells, while after low FSS preconditioning, ΔAP_cell_ and ΔAP_traction_ showed opposite and significant difference between Piezo1-inhibited and WT cells (Fig. 2K). These results suggest that Piezo1 inhibition disrupts cells’ parallel alignment partially under high FSS and perpendicular alignment completely under low FSS.

To investigate the temporal relationship between cell and traction alignment, time series analysis of AP_cell_ and AP_traction_ was performed (Fig. 2L, M). Under direct high FSS (Fig. 2L), Piezo1-inhibited cells exhibited a steady but attenuated increase in AP_cell_ and AP_traction_ over time. In contrast, after 60-min low FSS preconditioning (Fig. 2L), neither AP_cell_ nor AP_traction_ shows any significant trend over time, in stark contrast to the AP behaviors under direct high FSS condition. To seek if there is a functional causality between traction alignment and cell alignment, we performed Granger causality (GC) test, a statistical test used to determine if one time series predicts another. The details about the test is explained in the ref.^30^. Briefly, the test focuses on each causal direction, if AP_cell_ granger-causes (GCs) AP_traction_ or vice versa, produce a significance of causality as *P* values. To determine the global significance of each causal direction with harmonic mean P (HMP), the P value from several independent experiments per flow conditions were combined. We calculated the P values for pairs of AP timeseries under long-term 24 hours for direct high FSS and 60-min low FSS conditions for Piezo1 inhibited cells (Fig. 2N). The distribution of P values showed that in Piezo1 inhibited cells, the causal direction, ‘AP_traction_ GCs AP_cell_’ appeared to be more functional than the other causal direction for both direct high FSS and 60-min low FSS conditions (Fig. 2N). However, the Mann-Whitney test showed no significant difference between the directions. Thus, this analysis suggests that the inhibition of Piezo1 greatly attenuates not only the alignment behavior of the cells under flow but also the functional coupling between traction alignment and cell alignment. Together, these findings collectively suggest that Piezo1 partially mediates cell and traction alignment, coupling between cell and traction orientations, and traction magnitude under high FSS but largely uninfluential in translating low FSS to EC traction.

### TRPV4 governs endothelial rapid initial traction response to high and low FSS leading to slow traction relaxation and induced cell and traction alignment

To understand the role of TRPV4 in shear-mediated traction, we measured and analyzed the traction of HUVECs treated with TRPV4 siRNA in response to high FSS or 60-min low FSS preconditioning (Fig 3 A-D, S5 A-D). The knockdown efficiency was 89%, confirmed by qRCR (Fig. S2). In response to direct high FSS, TRPV4-inhibited HUVECs displayed a complete absence of the rapid traction increase seen in WT cells during the first 10 minutes (Fig. 3A, *orange curve*). Instead, a gradual, steady rise in traction began after 15 minutes and continued over time. This rising trend continued in the long term, reaching a global peak at ∼6 hour, followed by a slower relaxation that returned near baseline levels at 24 hour. This long-term rise is reminiscent of the second rise exhibited in control ECs exposed to 60-min low FSS (Fig. 2D, *gray*) or Piezo1-inhibited ECs exposed to direct FSS (Fig. 2B, *blue*), suggesting the idea of insufficient initial traction rise potentially associated with long-term traction elevation.

**Figure 3.**
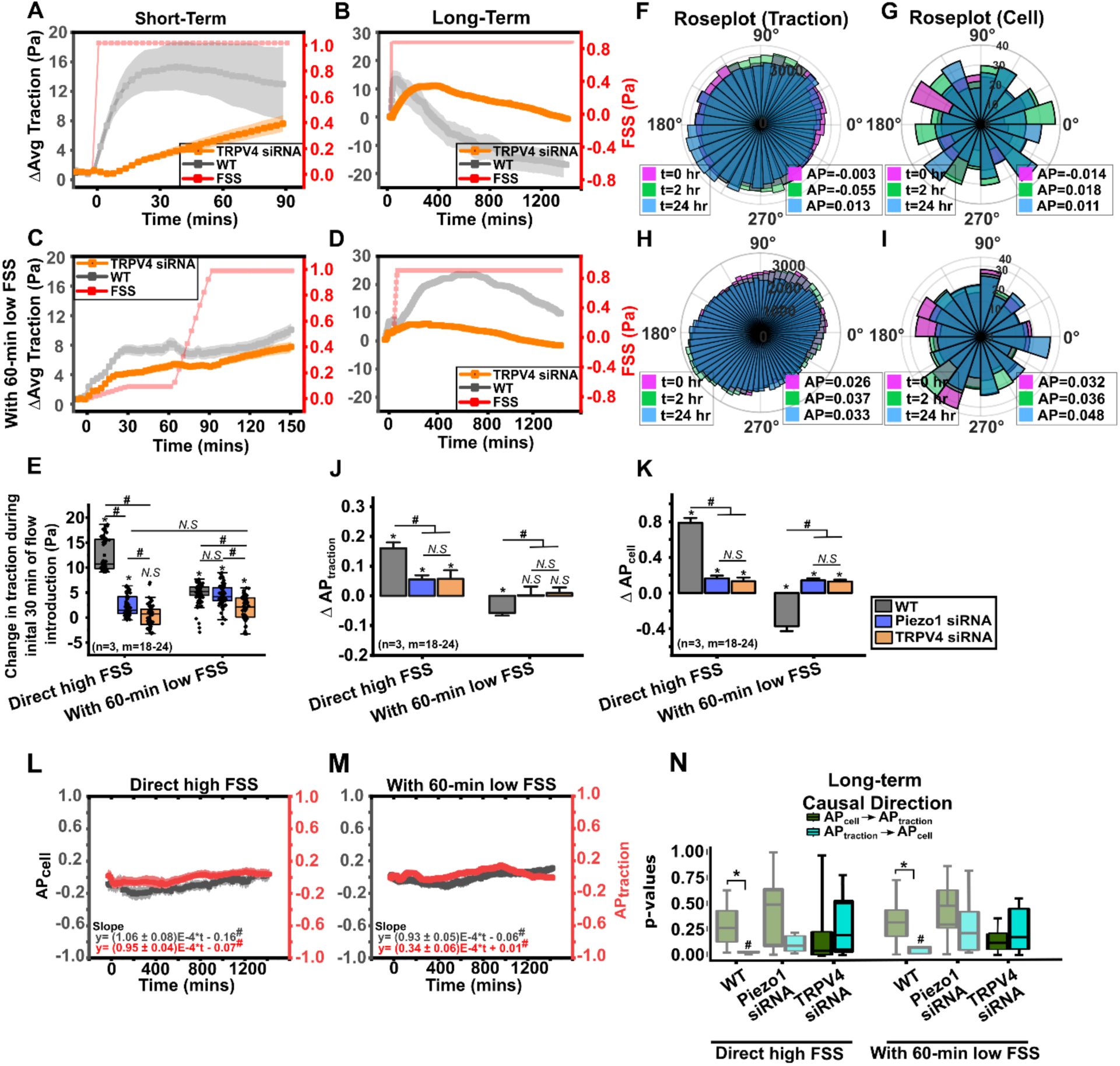
TRPV4 modulates endothelial traction response to onset of FSS. (A, B, C, D) Average traction of HUVECs silenced with TRPV4 siRNA (orange) for short-term (A, C) and long-term (B, D) under direct high FSS (A, B) and 60-min low FSS (C, D). Wildtype condition shown in (grey) and FSS in (red) (N=1, M=7-9). Error bar means ± SE. (E) A boxplot of early rise in traction within 30 min of FSS (N=3, M= 16-24). (F, H) Polar distribution of traction vectors at t=0 hr (magenta), t=2 hr (green) and t=24 hr (blue) for each condition. (G, I) Polar distribution of cell orientation at t=0 hr (magenta), t=2 hr (green) and t=24 hr (blue) for each condition. (J, K) Bar plot of changes in ΔAP_traction_ and ΔAP_cell_ for wild-type (*gray*), Piezo1siRNA (*blue*) and TRPV4siRNA (*orange*) under direct high FSS and 60-min low FSS conditions. (N=3, M= 18-24). (L, M) Time-series plot of alignment parameter, AP_cell_ (gray) and AP_traction_ (red) as a function of time from each representative time-lapse images recorded for direct high FSS (L), and 60-min low FSS (M). #*P*<0.05 tested with t-test for the slope of the line, compared against null hypothesis that the true slope is zero. N represents the number of independent experiments and M represents the position observed. \**P*<0.05 tested with one-sample t test, that is, compared against zero. # *P*<0.05 tested with one-way ANOVA with Tukey’s post hoc test. (N) Boxplots of P values from Granger causality (GC) tests for long-term imaging for all conditions with causal direction that “AP_cell_ GCs AP_traction_” (green) and “AP_traction_ GCs AP_cell_” (blue). #P < 0.05 tested with HMP-combined GC P values, *P < 0.05 tested with Mann–Whitney U test between the two-direction GC test P values. Sample sizes for each condition are N = 4, 10,12, 5, 10, 12 for direct high FSS (WT), direct high FSS (Piezo1 siRNA), direct high FSS (TRPV4 siRNA), 60-min low FSS (WT), 60-min low FSS (Piezo1 siRNA), 60-min low FSS (TRPV4 siRNA) respectively.

Under 60-min low FSS preconditioning, unlike Piezo1 inhibition case, TRPV4 inhibition caused near two-fold reduction in the initial traction increase compared to WT cells (Fig. 3C, E). This was followed by slow, constant rise in traction, which continued until t= ∼3 hours after which the traction decreased slowly (Fig. 3D). This relaxation is earlier and with lower maximum traction compared to WT cells, which suggests TRPV4’s role in sustained traction generation. Quantification of early traction changes within 30 minutes of FSS exposure revealed traction of TRPV4-inhibited cells reduced compared to Piezo1-inhibited cells as well for both flow conditions (Fig. 3E). These results demonstrate TRPV4’s critical role in early traction generation.

The traction orientation of TRPV4-inhibited cells stayed isotropic before and after the onset of direct high FSS (Fig. 3F, *pink and blue*). Similarly, under low FSS preconditioning, traction orientation exhibited no distinct change (Fig. 3H, *pink and blue*), which is in contrast with WT HUVECs’ traction alignment perpendicular to the flow under low FSS. Cell orientation (Fig. S6), depicted in rose plots (Fig. 3G, I), also remained isotropic regardless of the FSS condition in TRPV4-inhibited cells, showing no distinct alignment patterns after 24 hours under both direct high FSS (Fig. 3G) and low FSS preconditioning (Fig. 3I). AP for both traction and cell of the TRPV4-inhibited HUVECs also showed only slight increase for direct high FSS and no change for 60-min low FSS, similar to that of Piezo-1 inhibited HUVECs (Fig. 3J, K). This trend was also shown in the time series analysis with only small fluctuations of APs (Fig. 3L, M). GC analysis showed no significant functional causality between AP_cell_ and AP_traction_ (Fig. 3N). Together, these results suggest that TRPV4 mediates not only the immediate traction magnitude modulation upon the onset of FSS but also the coordination and coupling of alignment of the traction and the cells to the flow.

### Dual inhibition of Piezo1 and TRPV4 disrupts short-term traction response and alignment more completely

Studies with single ion channel inhibition revealed that neither Piezo1 nor TRPV4 alone could fully suppress shear-mediated traction forces. Piezo1 inhibition partially reduced the initial traction rise, while TRPV4 inhibition completely truncated the early traction response but allowed for a subsequent gradual increase during the short-term under direct high FSS (Figs. 2A and 3A). From these findings, we hypothesized that Piezo1 and TRPV4 may have complementary roles in mediating shear-induced traction. To test this hypothesis, we performed dual inhibition of both Piezo1 and TRPV4 and sought if this could more effectively downregulate shear-mediated traction in HUVECs (Fig. 4). In the dual knockdown group, the knockdown efficiency of Piezo1 was 88.9% and TRPV4 was 85.4% (Fig. S1, S2). In response to direct high FSS, dual-inhibited cells exhibited a small initial increase in traction, after which it stayed constant at ∼3 Pa for the first 90 minutes (Fig. 4A, E, Fig. S7A). This finding suggests that Piezo1 and TRPV4 are key, complementary mechanosensitive ion channels responsible for generating the rapid traction forces necessary for ECs to respond to shear stress in the short-term.

**Figure 4.**
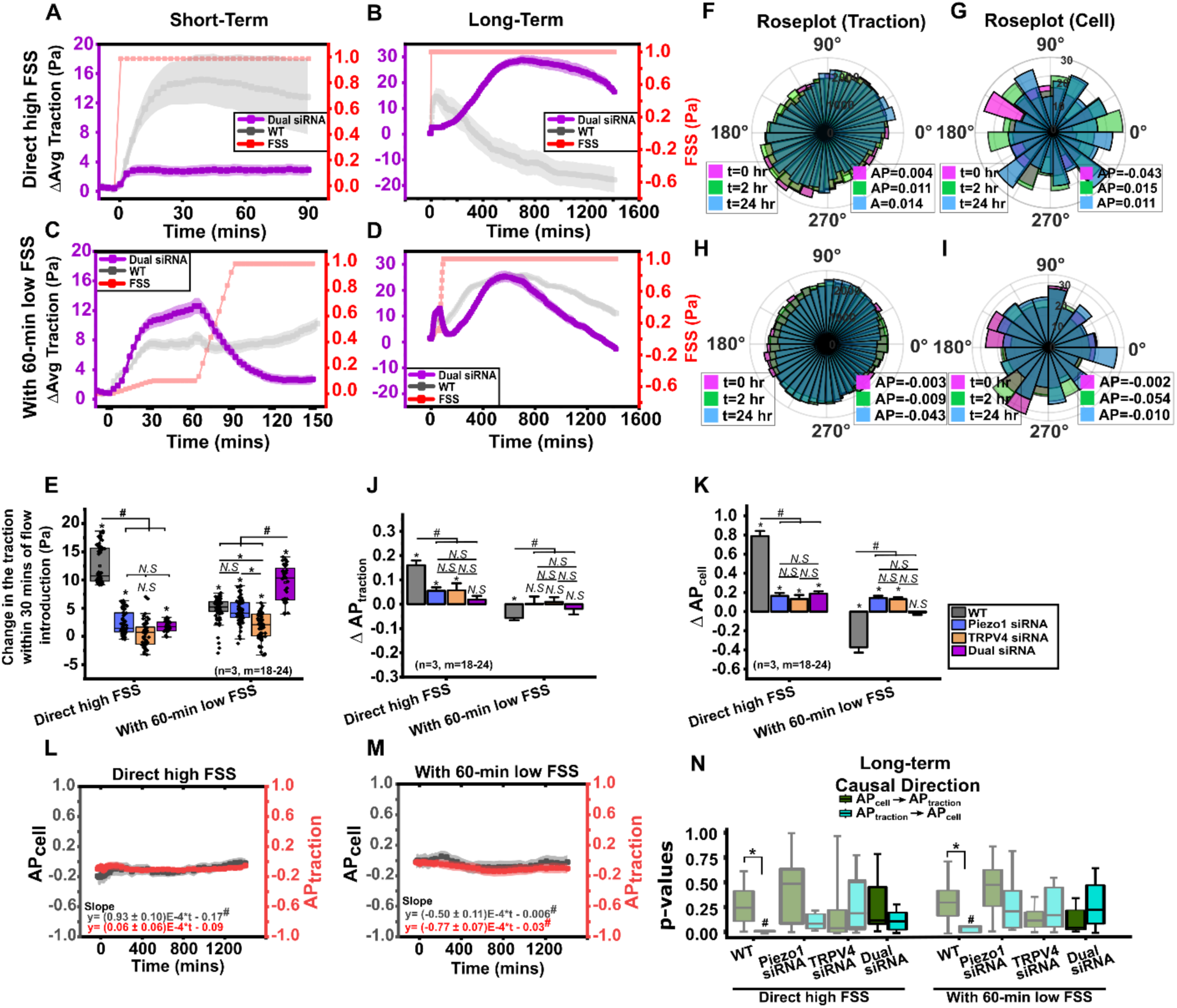
Piezo1 and TRPV4 inhibition together more thoroughly inhibits the early traction response further leading to secondary traction rise in long-term in response to high FSS. (A, B, C, D) Average traction of HUVECs silenced with piezo1 and TRPV4 siRNA (purple) for short-term (A, C) and long-term (B, D) under direct high FSS (A, B) and 60-min low FSS (C, D). Wildtype condition shown in (grey) (N=1, M=7-9). Error bar means ± SE. (E) A boxplot of early rise in traction within 30 min of FSS (N=3, M= 16-24). (F, H) Polar distribution of traction vectors at t=0 hr (magenta), t=2 hr (green) and t=24 hr (blue) for each condition. (G, I) Polar distribution of cell orientation at t=0 hr (magenta), t=2 hr (green) and t=24 hr (blue) for each condition. (J, K) Bar plot of changes in ΔAPtraction and ΔAP_cell_ for wild-type (*gray*), Piezo1siRNA (*blue*), TRPV4siRNA (*orange*), and dualsiRNA (*purple*) under direct high FSS and 60-min low FSS conditions. (N=3, M= 18-24). (L, M) Time-series plot of alignment parameter, AP_cell_ (gray) and AP_traction_ (red) as a function of time from each representative time-lapse images recorded for direct high FSS (L), and 60-min low FSS (M). #*P*<0.05 tested with t-test for the slope of the line, compared against null hypothesis that the true slope is zero. N represents the number of independent experiments and M represents the position observed. \**P*<0.05 tested with one-sample t test, that is, compared against zero. # *P*<0.05 tested with one-way ANOVA with Tukey’s post hoc test. (N) Boxplots of P values from Granger causality (GC) tests for long-term imaging for all conditions with causal direction that “AP_cell_ GCs AP_traction_” (green) and “AP_traction_ GCs AP_cell_” (blue). #P < 0.05 tested with HMP-combined GC P values, *P < 0.05 tested with Mann–Whitney U test between the two-direction GC test P values. Sample sizes for each condition are N = 4, 10, 12, 10 5, 10, 12, 10 for direct high FSS (WT), direct high FSS (Piezo1 siRNA), direct high FSS (TRPV4 siRNA), direct high FSS (Dual siRNA), 60-min low FSS (WT), 60-min low FSS (Piezo1 siRNA), 60-min low FSS (TRPV4 siRNA), 60-min low FSS (Dual siRNA) respectively.

However, after a short-term, this low, steady phase was followed by a gradual increase in traction over the next several hours, eventually reaching a global peak at 12 hours (Fig. 4B, S7B) that exceeded the maximum observed in WT cells. The subsequent relaxation phase was slow, and traction did not return to baseline levels within 24 hours (Fig. 4B).

Conversely, to our surprise, the average traction of dual-inhibited cells showed an increase 1.5 times larger than that of WT cells in response to low FSS (Fig. 4C, E, S7 C). However, as FSS was ramped from 0.1 Pa to 1 Pa, traction began to decline, returning close to baseline levels shortly after the transition to high FSS. This immediate inverse behavior suggests that upon double-inhibition, calcium-independent pathways become dominant and are more directly sensitive to the high FSS. In the long term, this reduced traction started to rise again (Fig. 4D, S7D), which is similar to the secondary elevation shown in WT cells.

The traction orientation of dual-inhibited cells showed insignificant change over 24 hours under direct high FSS (Fig. 4F, J), so did traction under low FSS preconditioning (Fig. 4H, J). Likewise, the cell orientation distributions (Fig. S8) remained largely isotropic, with no significant changes under both direct high FSS (Fig. 4G, K) and low FSS preconditioning (Fig. 4I, K), which resembles to that of TRPV4-inhibited condition. AP time series also showed minimal changes over time (Fig. 4L, M) with no functional causality between AP_traction_ and AP_cell_ (Fig. 4N), similar to Piezo1- and TRPV4- inhibited HUVECs. These results collectively suggest that dual inhibition of Piezo1 and TRPV4 mediates both immediate and sustained traction magnitude modulation under FSS and impairs the alignment of traction and cell to the flow.

### Integrin Signaling is Dispensable for Secondary Traction Rise, but May Modulate Short-Term TRPV4 Responses

All FSS-TM experiments with ion channel inhibition under high or low FSS showed secondary traction rise in the long term where traction gradually increased over several hours, reaching a peak before slowly relaxing. This secondary rise was not shown only in WT HUVECs under direct high FSS. To test whether this secondary rise is from integrin signaling, we performed FSS-TM experiments with the inhibition of focal adhesion kinase (FAK), a key integrin signaling molecule. Upon inhibiting FAK, under direct high FSS, the short-term traction response exhibited a noticeable decrease in overall traction but was not completely eliminated (Fig. S9A, E). However, this was followed by the secondary rise in traction (Fig. S9B). Likewise, under 60-min low FSS, a secondary rise almost identical to those observed in WT cells was present (Fig. S9C, D). This data suggests that the secondary traction rise is independent of FAK signaling, indicating the involvement of an alternative pathway. Interestingly, the short-term traction behavior of FAK-inhibited cells under direct high shear closely resembled that observed in TRPV4-inhibited cells (Fig. S9A vs. Fig. 3A). Moreover, the inverse relationship between increased FSS and decreased traction was also observed in FAK-inhibited cells (Fig. S9D). Thus, while our results suggest that the secondary traction rise is not mediated by FAK, we cannot rule out the possibility that calcium ion channels regulate traction through integrin activation.

## Discussion

Our study highlights the essential roles of Piezo1 and TRPV4 mechanosensitive ion channels in EC responses to FSS, specifically in traction magnitude modulation and alignment under varying flow conditions. Overall, the inhibition of these ion channels resulted in reduced traction behavior when subjected to high FSS and transient low FSS in comparison to the WT condition. These initial reduced traction with the onset of the FSS leads to long-term traction elevation further affecting the traction relaxation behavior. Moreover, these early traction response from both direct high FSS and transient low FSS leads to isotropic distribution of the traction vectors further affecting the cell orientation toward no significant alignment to flow direction. The findings provide critical insights into how these ion channels function independently and collaboratively to regulate endothelial mechanotransduction.

Our study reveals that Piezo1 shows sensitivity to high FSS but not to transient low FSS. Under high FSS, traction rise was greatly diminished, whereas under transient low FSS, traction response was comparable to that of WT cells. This observation aligns with findings that Piezo1 is activated by high FSS and regulates calcium activity^12,36^. As calcium activity controls the focal adhesion assembly and disassembly^11,37^, the reduced traction responses under direct high FSS might be due to inhibited calcium-dependent FA remodeling. Particularly, we speculate that reduced calcium entry by Piezo1 inhibition could have reduced FA assembly transiently then blocked the disassembly of FAs, which might be responsible for the secondary rise.

Conversely, the indistinct traction response of Piezo-inhibited cells to transient low FSS compared to WT HUVECS may be attributed to specific ECM protein’s effect in FSS range sensing^38^. Piezo1-induced calcium influx has been shown to depend on the magnitude of the FSS and selectively activate specific integrin heterodimers^12^. Notably, under low FSS conditions, cells on collagen-coated substrates exhibited increased calcium signaling through Piezo1 activation, mediated by interactions with α_v_ integrins. In contrast, cells on fibronectin-coated substrates showed a delayed and reduced calcium response, indicating that different extracellular matrix components modulate Piezo1 mechanosensitivity in an FSS magnitude-dependent manner^12^. This implies that under low FSS, Piezo1 plays a minimal role in the presence of fibronectin, and its inhibition does not significantly alter traction forces. Instead, the integrin-fibronectin interaction likely drives traction through Piezo1-independent pathways, explaining why Piezo1-inhibited cells exhibit similar traction responses to wild-type cells under low FSS conditions.

Our data show that TRPV4 inhibition completely hampered the short-term traction increase against direct high FSS during the first 15 minutes. This observation is consistent with findings with the calcium influx suppression from TRPV4-inhibited ECs in response to FSS^39^. Additionally, TRPV4 inhibition not only suppresses calcium activity but also disrupts β-catenin-dependent signaling pathways, as TRPV4 forms a complex with β-catenin^40^. These complexes reside both at cell-cell junctions and basal membrane in static culture, but the application of the flow separates TRPV4 from β-catenin and facilitate FAK and integrin signaling^40^. Therefore, TRPV4 inhibition may impair integrin activation, leading to reduced traction forces under high FSS. Similarly, the diminished traction response observed in TRPV4-inhibited HUVECs under low FSS preconditioning may be partially attributed to the disruption of TRPV4-β-catenin interactions and FAK signaling. Furthermore, TRPV4 inhibition may also affect Piezo1-mediated pathways. Studies suggest that TRPV4 activation occurs downstream of Piezo1^7^, indicating that both channels may converge on shared mechanotransduction pathways.

Interestingly, the slow and steady traction increase observed after 15 minutes under TRPV4 inhibition may indicate compensatory activation of alternative mechanosensors, such as glycocalyx^41^, other ion channels^42^, and GPCRs^43^, followed by their downstream pathways. Alternative explanation could involve the side effect of integrin activity inhibition where FAK Y397 phosphorylation inhibition leads to inhibition of focal adhesion turnover and growth of FAs^44^. This compensation highlights the adaptability of ECs to sustain contractility in the absence of TRPV4. These findings suggest that TRPV4 plays a central role in early endothelial remodeling.

Our dual inhibition against both Piezo1 and TRPV4 showed a nearly 5-fold decrease in overall traction compared to WT under direct high FSS. Piezo1 and TRPV4 together are critical for almost 80% of the traction response to direct high FSS. Piezo1 is known for elevating inflammation^8^, while TRPV4 leads to vasodilation^19^; thus, blocking both may suppress two independent calcium-dependent pathways^45^, reducing traction forces.

On the contrary, our dual inhibition results show a one-fold increase in traction compared to WT conditions in response to low shear preconditioned flow. This traction increase may be attributed to calcium-independent mechanosensors such as Kir^46^ or GPCRs^43^, or calcium-independent pathways such as phospholipase C-β1 via GPCRs, protein kinases like PKC, and MAP kinases^45^, though their exact activation mechanisms and downstream pathways require further investigation. These pathways are associated with vasodilation by producing NO and PGI2, inflammation, and immune response by controlling adhesion molecules and cytokines^47^. It is possible that low shear activates a series of calcium-independent downstream pathways involved in vasodilation, inflammation, and immune response, resulting in increased traction. Mapping these pathways and their involvement in traction generation requires further exploration.

In conclusion, our data demonstrate that TRPV4 and Piezo1 are critical mechanosensitive ion channels that regulate endothelial traction force generation, alignment, and adaptation, particularly under high FSS. TRPV4 emerges as the dominant regulator, orchestrating both the initial traction rise and long-term relaxation, whereas Piezo1 contributes primarily to the early response partially under high FSS. While many mechanistic details remain to be elucidated, our findings suggest that ion channels serve as the first line of FSS sensing, with TRPV4 playing a central role in translating shear stress magnitude into endothelial mechanotransduction. These insights provide a deeper understanding of endothelial adaptation to flow and its implications for vascular pathophysiology.

## Acknowledgement

We thank Dr. Feng Zhao (Texas A&M University) for sharing early passage HUVECs and helping us with HUVEC culture protocols, Sang Joon Ahn (UIC, Chicago), Yeonwoo Rho (MTU) and Paul Evans (Queen Mary University London, UK) for helpful discussions, and Aimee Marceau from Genomic Sequencing Lab (MTU) for assisting with qPCR facility.

## Author Contributions

M.K.C. and S.J.H. conceived and designed research; M.K.C., M.Y. performed experiments; M.K.C., M.Y., M.R., and S.J.H. analyzed data; M.K.C. and S.J.H. interpreted results of experiments; M.K.C. prepared figures; M.K.C. drafted manuscript; S.J.H. edited and revised manuscript; S.J.H. approved final version of manuscript.

## Supplemental Figures

**Fig S1.**
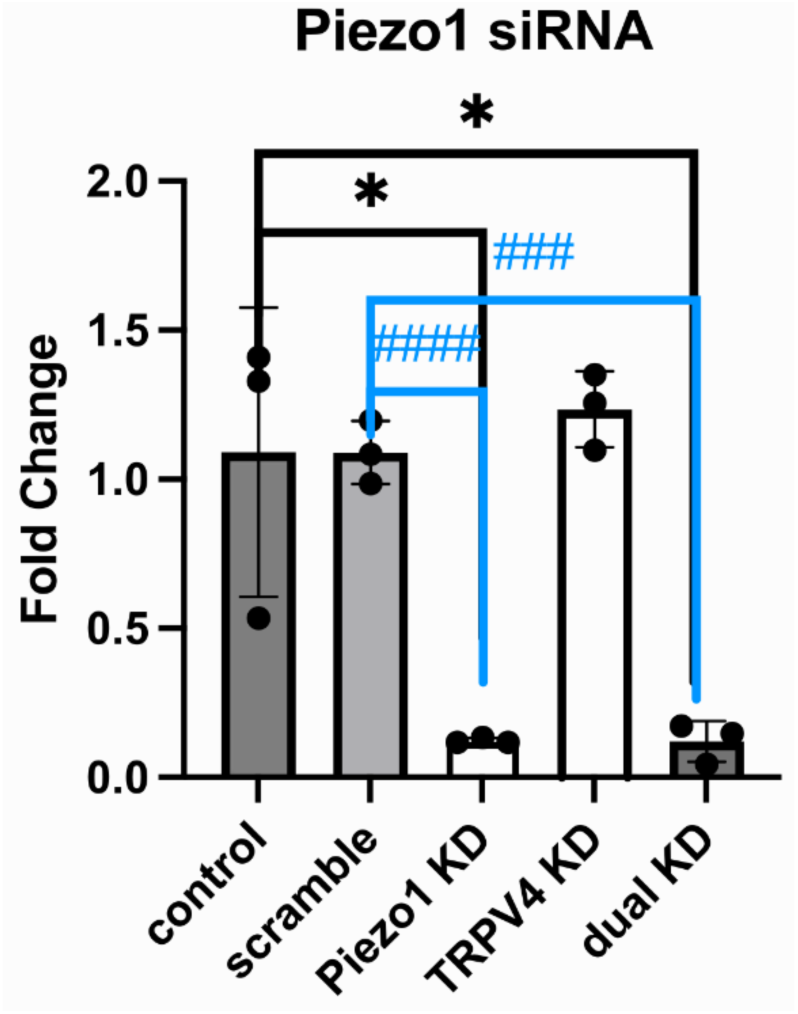
As shown in the figure, there was no change in the expression level of piezo1 in the control group compared to the scramble si-RNA negative control group. Compared to the control group, the expression levels of piezo1 in the piezo1 knockdown group and the dual knockdown group were significantly reduced, while the expression level of piezo1 in the trpv4 knockdown group remained unchanged. Similarly, compared to the scramble si-RNA negative control group, the expression levels of piezo1 in the piezo1 knockdown group and the dual knockdown group were significantly reduced, with no change observed in the piezo1 expression level in the trpv4 knockdown group.

**Fig S2.**
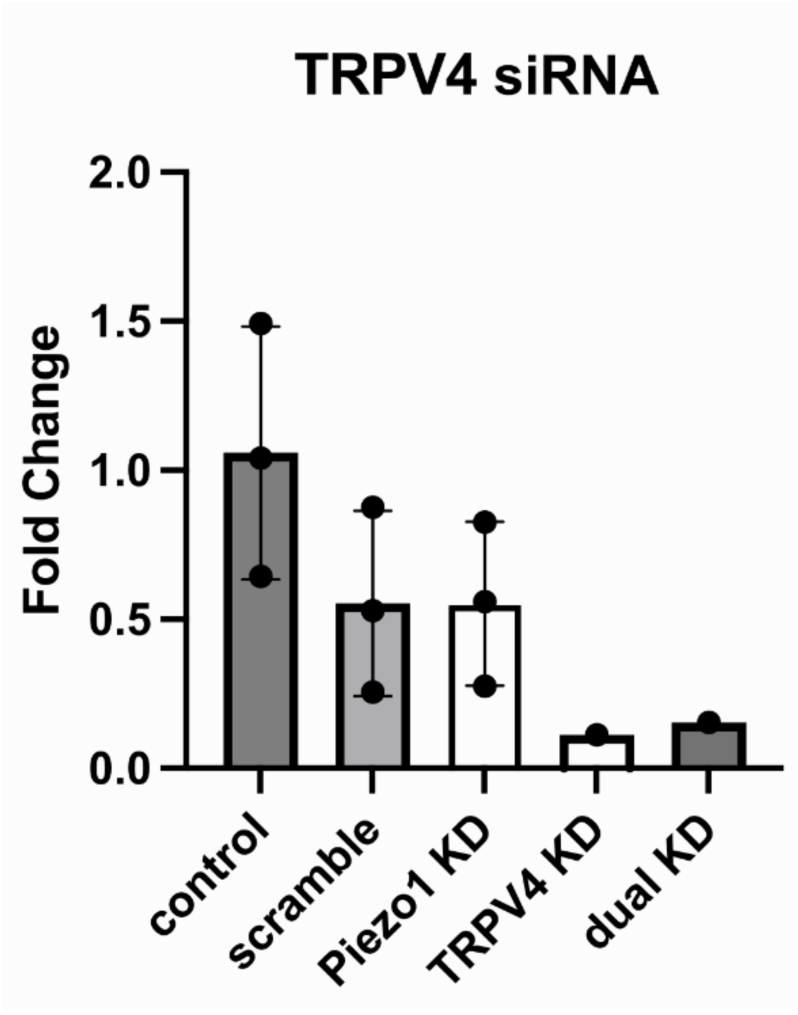
As shown in the figure, the expression of the TRPV4 gene in both the TRPV4 KD group and the Dual KD group is significantly reduced, confirming successful knockdown. For the TRPV4 KD group, the Ct values are 34.004, Undetermined, and Undetermined across the three wells. Similarly, in the Dual KD group, the Ct values are 33.927, Undetermined, and 35.402. The presence of high Ct values and several Undetermined wells in both groups suggests that the TRPV4 gene expression was knocked down effectively by the TRPV4 siRNA, reducing the expression to levels that are near or below the detection threshold of the system. The original qPCR data is attached in the file.

**Fig S3.**
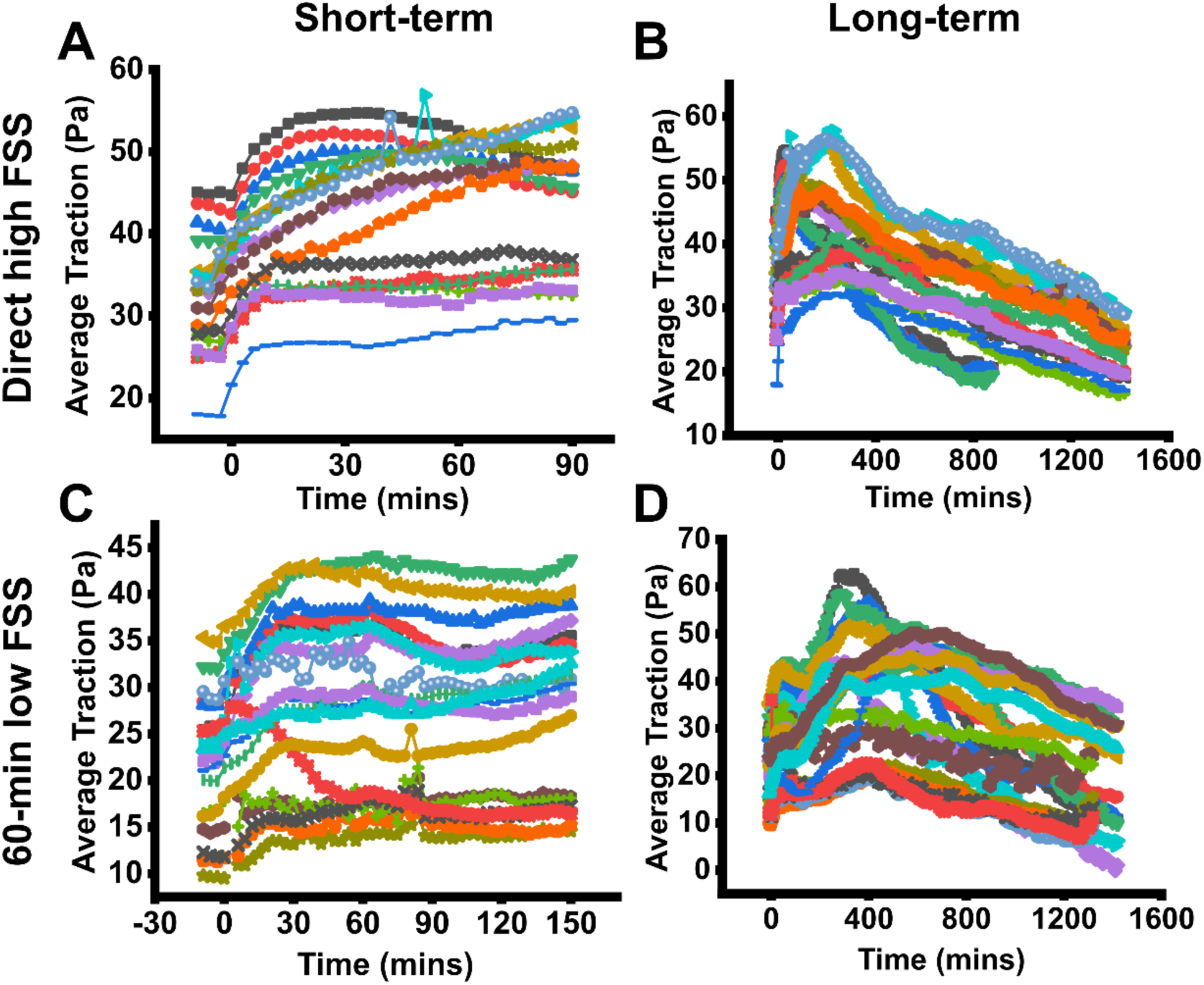
Average traction magnitude as a function of time for each individual region of interests, co-plotted with FSS profiles on Piezo1 inhibited ECs. (A, C) Short-term traction response for direct high FSS and 60-min low FSS (M=3, N=16-24). (B, D) Long-term traction response for direct high FSS and 60-min low FSS (M=3, N=16-24). Here, M represents the number of flow experiments and N represents the number of regions of interest observed in each experiment.

**Fig S4.**
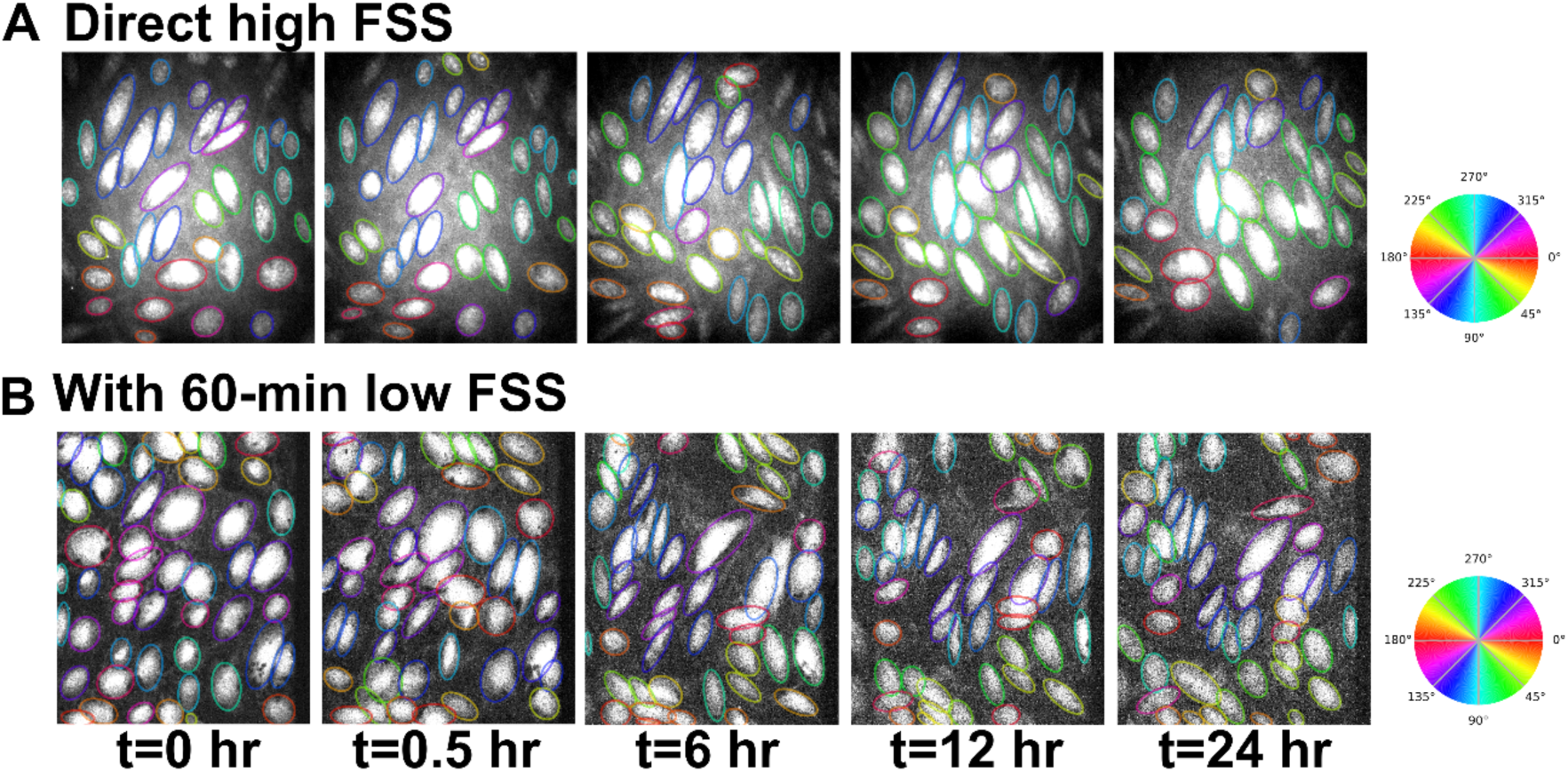
Predicted segmentation for direct high FSS (A) and 60-min low FSS (B) in Piezo1 inhibited ECs for time-points t= 0, 0.5, 6, 12, and 24 hours with angular color map showing the segmented cells with respective angular distribution color coding in the field of view.

**Fig S5.**
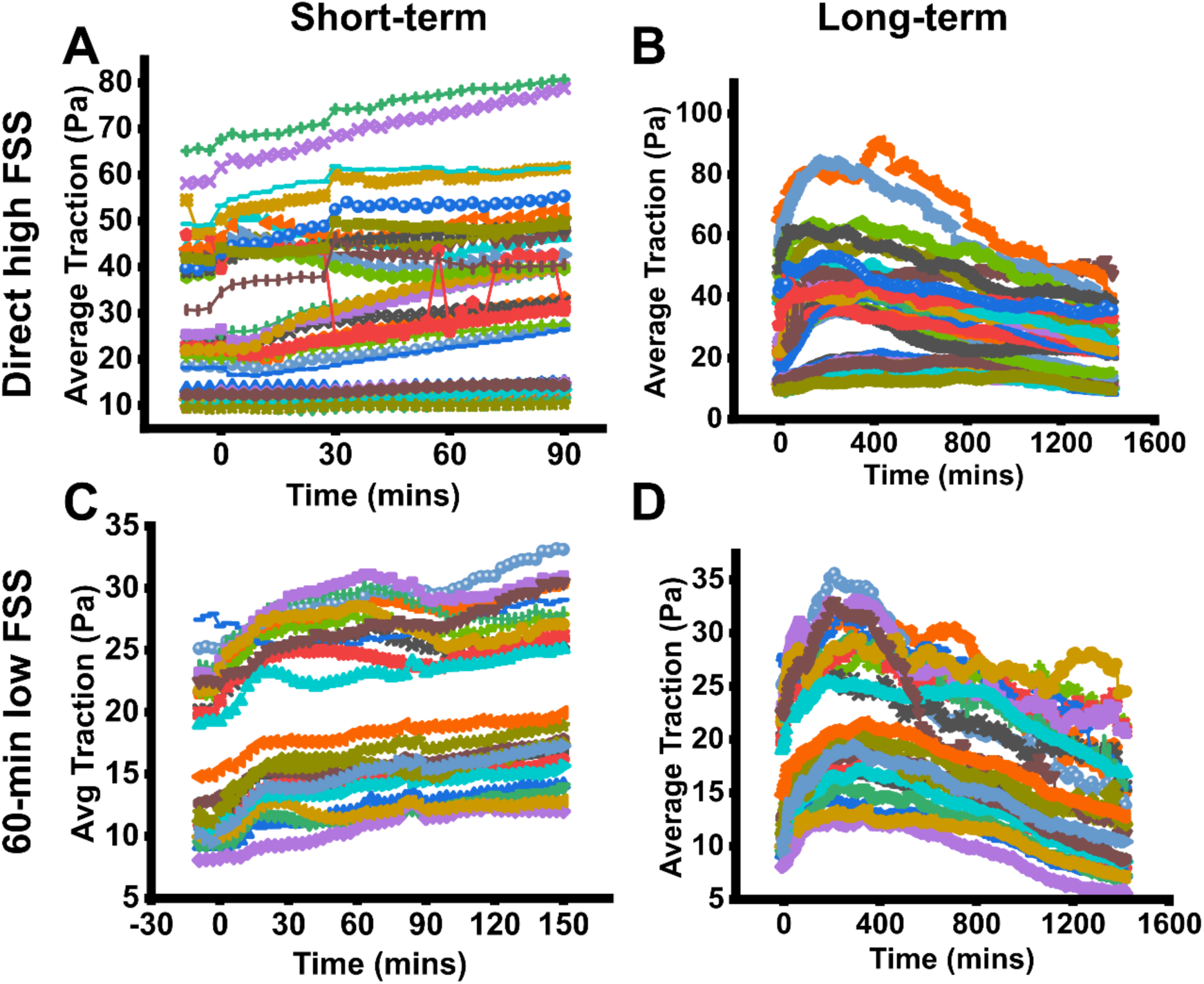
Average traction magnitude as a function of time for each individual region of interests, co-plotted with FSS profiles on TRPV4 inhibited ECs. (A, C) Short-term traction response for direct high FSS and 60-min low FSS (M=3, N=16-24). (B, D) Long-term traction response for direct high FSS and 60-min low FSS (M=3, N=16-24). Here, M represents the number of flow experiments and N represents the number of regions of interest observed in each experiment.

**Fig S6.**
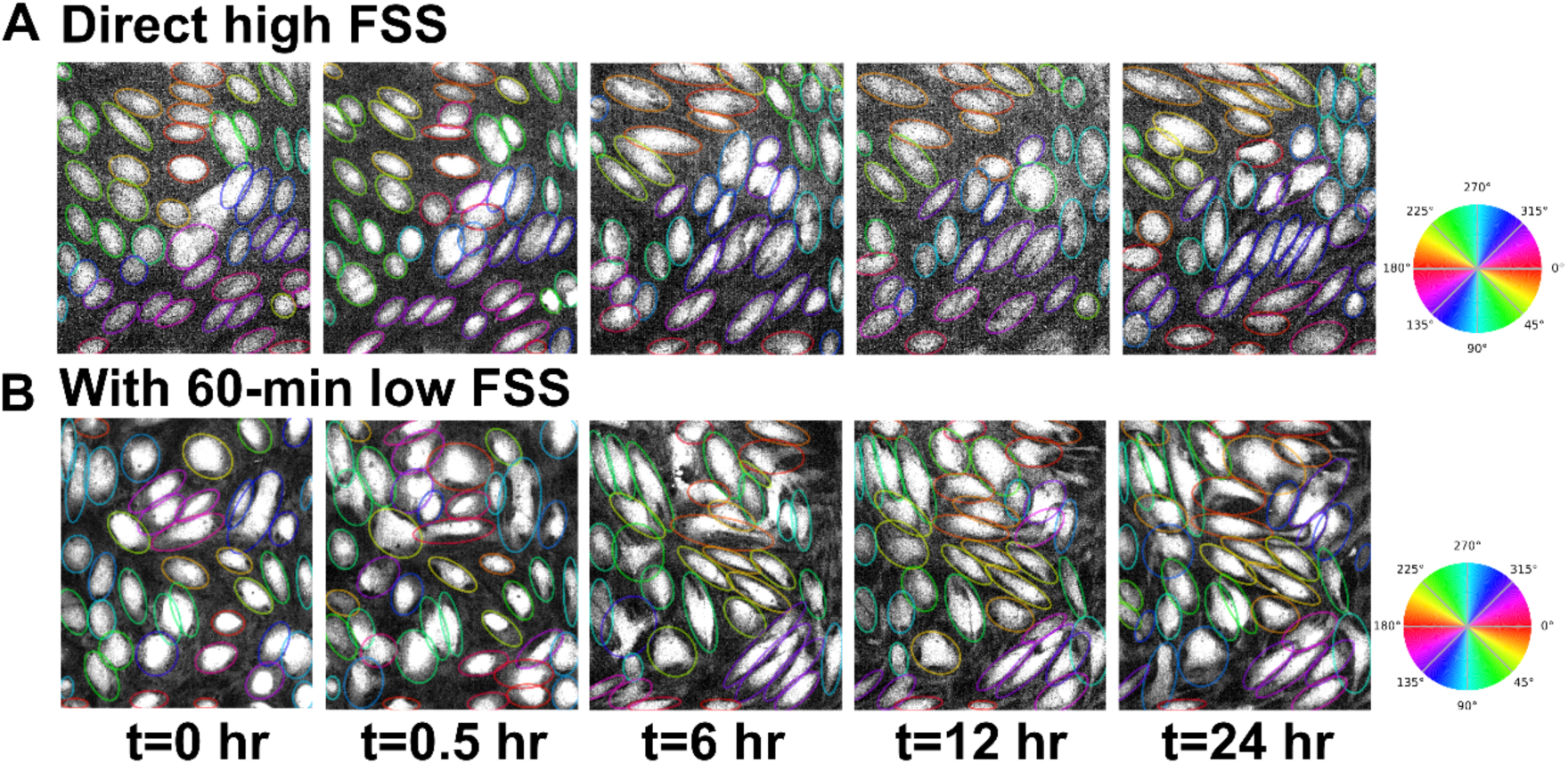
Predicted segmentation for direct high FSS (A) and 60-min low FSS (B) in TRPV4 inhibited ECs for time-points t= 0, 0.5, 6, 12, and 24 hours with angular color map showing the segmented cells with respective angular distribution color coding in the field of view.

**Fig S7.**
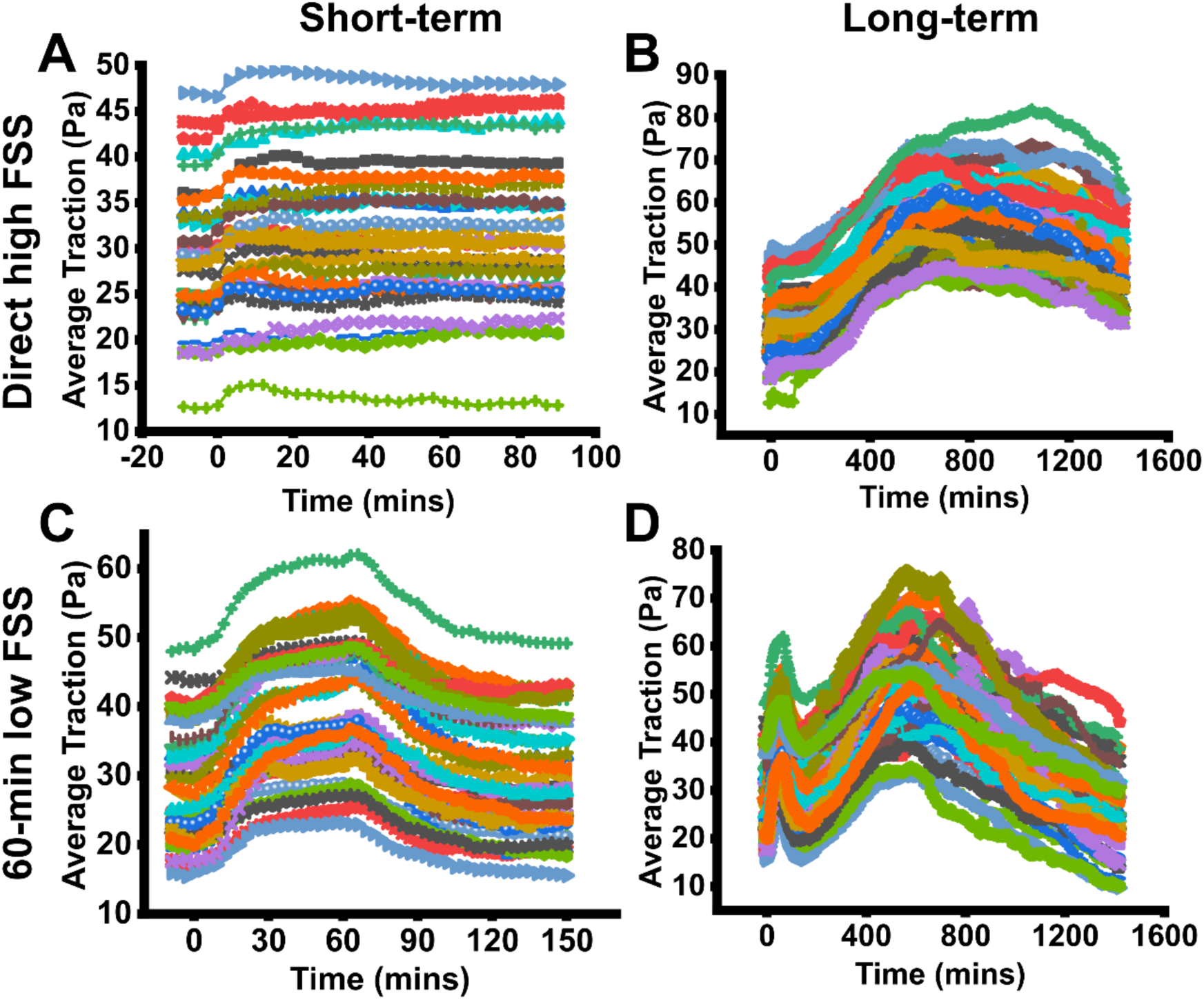
Average traction magnitude as a function of time for each individual region of interests, co-plotted with FSS profiles on dual inhibited ECs. (A, C) Short-term traction response for direct high FSS and 60-min low FSS (M=3, N=16-24). (B, D) Long-term traction response for direct high FSS and 60-min low FSS (M=3, N=16-24). Here, M represents the number of flow experiments and N represents the number of regions of interest observed in each experiment.

**Fig S8.**
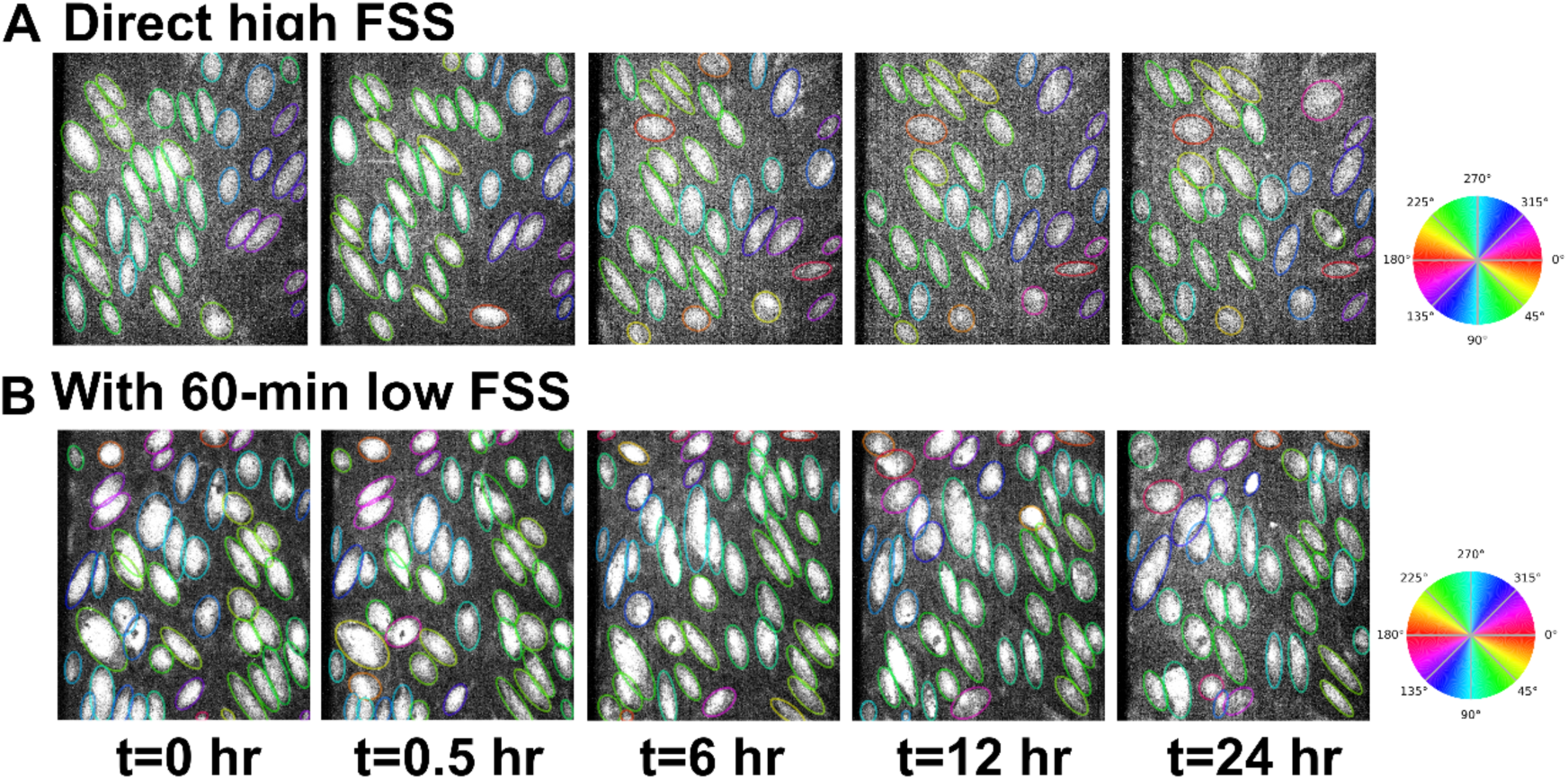
Predicted segmentation for direct high FSS (A) and 60-min low FSS (B) in dual inhibited ECs for time-points t= 0, 0.5, 6, 12, and 24 hours with angular color map showing the segmented cells with respective angular distribution color coding in the field of view.

**Fig S9.**
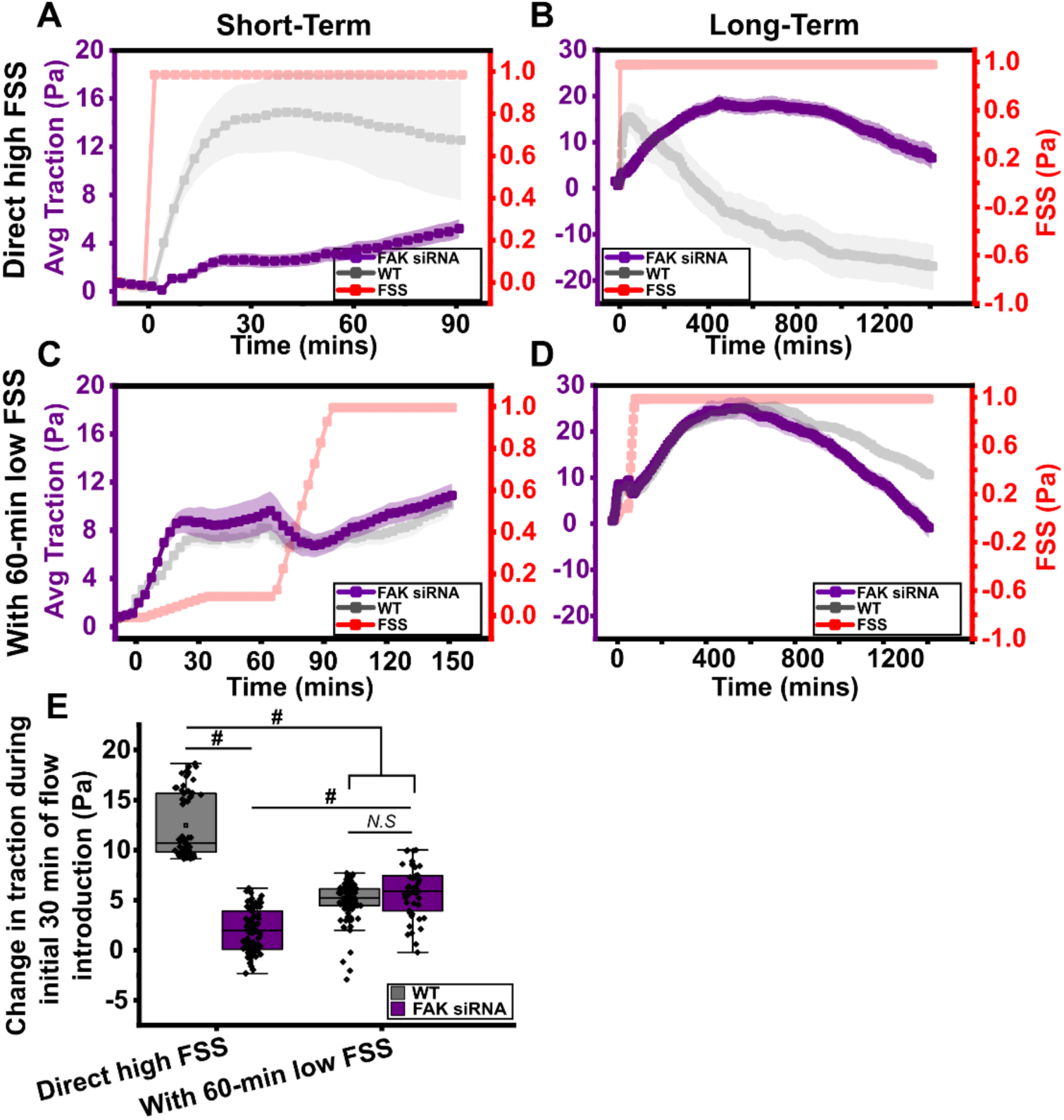
(A, B, C, D) Average traction of HUVECs silenced with FAK siRNA (violet) for short-term (A, C) and long-term (B, D) under direct high FSS (A, B) and 60-min low FSS (C, D). Wildtype condition shown in (grey) (N=1, M=7-9). Error bar means ± SE. (E) A boxplot of early rise in traction within 30 min of FSS (N=3, M= 16-24).

## References

1. Heo K-S, Fujiwara K, Abe J-i. Shear stress and atherosclerosis. Molecules and cells. 2014;37:435.

2. Lekka M, Gnanachandran K, Kubiak A, Zieliński T, Zemła J. Traction force microscopy – Measuring the forces exerted by cells. Micron. 2021;150:103138. doi: 10.1016/j.micron.2021.103138

3. Albarrán-Juárez J, Iring A, Wang S, Joseph S, Grimm M, Strilic B, Wettschureck N, Althoff TF, Offermanns S. Piezo1 and Gq/G11 promote endothelial inflammation depending on flow pattern and integrin activation. Journal of Experimental Medicine. 2018;215:2655–2672. doi: 10.1084/jem.20180483

4. Hahn C, Schwartz MA. Mechanotransduction in vascular physiology and atherogenesis. Nature Reviews Molecular Cell Biology. 2009;10:53–62. doi: 10.1038/nrm2596

5. Haga JH, Li Y-SJ, Chien S. Molecular basis of the effects of mechanical stretch on vascular smooth muscle cells. Journal of biomechanics. 2007;40:947–960.

6. Shah V, Patel S, Shah J. Emerging role of Piezo ion channels in cardiovascular development. Developmental Dynamics. 2022;251:276–286. doi: 10.1002/dvdy.401

7. Swain SM, Liddle RA. Piezo1 acts upstream of TRPV4 to induce pathological changes in endothelial cells due to shear stress. Journal of Biological Chemistry. 2021;296. doi: 10.1074/jbc.RA120.015059

8. Yang Y, Wang D, Zhang C, Yang W, Li C, Gao Z, Pei K, Li Y. Piezo1 mediates endothelial atherogenic inflammatory responses via regulation of YAP/TAZ activation. Human Cell. 2022;35:51–62. doi: 10.1007/s13577-021-00600-5

9. Li J, Hou B, Tumova S, Muraki K, Bruns A, Ludlow MJ, Sedo A, Hyman AJ, McKeown L, Young RS, et al. Piezo1 integration of vascular architecture with physiological force. Nature. 2014;515:279–282. doi: 10.1038/nature13701

10. Coste B, Mathur J, Schmidt M, Earley TJ, Ranade S, Petrus MJ, Dubin AE, Patapoutian A. Piezo1 and Piezo2 are essential components of distinct mechanically activated cation channels. Science. 2010;330:55–60. doi: 10.1126/science.1193270

11. D’Souza RS, Lim JY, Turgut A, Servage K, Zhang J, Orth K, Sosale NG, Lazzara MJ, Allegood J, Casanova JE. Calcium-stimulated disassembly of focal adhesions mediated by an ORP3/IQSec1 complex. eLife. 2020;9:e54113. doi: 10.7554/eLife.54113

12. Lai A, Thurgood P, Cox CD, Chheang C, Peter K, Jaworowski A, Khoshmanesh K, Baratchi S. Piezo1 Response to Shear Stress Is Controlled by the Components of the Extracellular Matrix. ACS Applied Materials & Interfaces. 2022;14:40559–40568.

13. Li J, Hou B, Tumova S, Muraki K, Bruns A, Ludlow MJ, Sedo A, Hyman AJ, McKeown L, Young RS. Piezo1 integration of vascular architecture with physiological force. Nature. 2014;515:279–282.

14. Chistiakov DA, Orekhov AN, Bobryshev YV. Effects of shear stress on endothelial cells: go with the flow. Acta Physiologica. 2017;219:382–408. doi: 10.1111/apha.12725

15. Zheng Q, Zou Y, Teng P, Chen Z, Wu Y, Dai X, Li X, Hu Z, Wu S, Xu Y. Mechanosensitive channel PIEZO1 senses shear force to induce KLF2/4 expression via CaMKII/MEKK3/ERK5 axis in endothelial cells. Cells. 2022;11:2191.

16. Wang S, Chennupati R, Kaur H, Iring A, Wettschureck N, Offermanns S. Endothelial cation channel PIEZO1 controls blood pressure by mediating flow-induced ATP release. The Journal of Clinical Investigation. 2016;126:4527–4536. doi: 10.1172/JCI87343

17. Köhler R, Hoyer J. Role of TRPV4 in the mechanotransduction of shear stress in endothelial cells. TRP ion channel function in sensory transduction and cellular signaling cascades. 2007.

18. Köhler R, Heyken W-T, Heinau P, Schubert R, Si H, Kacik M, Busch C, Grgic I, Maier T, Hoyer J. Evidence for a Functional Role of Endothelial Transient Receptor Potential V4 in Shear Stress–Induced Vasodilatation. *Arteriosclerosis*, Thrombosis, and Vascular Biology. 2006;26:1495–1502. doi: doi:10.1161/01.ATV.0000225698.36212.6a

19. Mendoza SA, Fang J, Gutterman DD, Wilcox DA, Bubolz AH, Li R, Suzuki M, Zhang DX. TRPV4-mediated endothelial Ca2+ influx and vasodilation in response to shear stress. American Journal of Physiology-Heart and Circulatory Physiology. 2010;298:H466–H476.

20. Thodeti CK, Matthews B, Ravi A, Mammoto A, Ghosh K, Bracha AL, Ingber DE. TRPV4 Channels Mediate Cyclic Strain–Induced Endothelial Cell Reorientation Through Integrin-to-Integrin Signaling. Circulation Research. 2009;104:1123–1130. doi: 10.1161/CIRCRESAHA.108.192930

21. Liu L, Guo M, Lv X, Wang Z, Yang J, Li Y, Yu F, Wen X, Feng L, Zhou T. Role of Transient Receptor Potential Vanilloid 4 in Vascular Function. Frontiers in Molecular Biosciences. 2021;8. doi: 10.3389/fmolb.2021.677661

22. Lewis AH, Grandl J. Mechanical sensitivity of Piezo1 ion channels can be tuned by cellular membrane tension. eLife. 2015;4:e12088. doi: 10.7554/eLife.12088

23. Chandurkar MK, Mittal N, Royer-Weeden SP, Lehmann SD, Michels EB, Haarman SE, Severance SA, Rho Y, Han SJ. Transient Low Shear Stress Preconditioning Influences Long-term Endothelial Traction and Alignment under High Shear Flow. Am J Physiol Heart Circ Physiol. 2024. doi: 10.1152/ajpheart.00067.2024

24. Mittal N, Michels EB, Massey AE, Qiu Y, Royer-Weeden SP, Smith BR, Cartagena-Rivera AX, Han SJ. Myosin-independent stiffness sensing by fibroblasts is regulated by the viscoelasticity of flowing actin. Communications Materials. 2024;5:6. doi: 10.1038/s43246-024-00444-0

25. Mittal N, Michels E, Pakenas K, Royer S, Han S. Polymerizing actin regulates myosin-independent mechanosensing by modulating actin elasticity and flow fluctuation. 2023.

26. Lane WO, Jantzen AE, Carlon TA, Jamiolkowski RM, Grenet JE, Ley MM, Haseltine JM, Galinat LJ, Lin F-H, Allen JD. Parallel-plate flow chamber and continuous flow circuit to evaluate endothelial progenitor cells under laminar flow shear stress. JoVE (Journal of Visualized Experiments). 2012:e3349.

27. Baeyens N, Nicoli S, Coon BG, Ross TD, Van den Dries K, Han J, Lauridsen HM, Mejean CO, Eichmann A, Thomas J-L, et al. Vascular remodeling is governed by a VEGFR3-dependent fluid shear stress set point. eLife. 2015;4:e04645. doi: 10.7554/eLife.04645

28. Papaioannou TG, Stefanadis C. Vascular wall shear stress: basic principles and methods. Hellenic J Cardiol. 2005;46:9–15.

29. Yin W, Shanmugavelayudam SK, Rubenstein DA. The effect of physiologically relevant dynamic shear stress on platelet and endothelial cell activation. Thrombosis research. 2011;127:235–241.

30. Chandurkar MK, Mittal N, Royer-Weeden SP, Lehmann SD, Michels EB, Haarman SE, Severance SA, Rho Y, Han SJ. Transient low shear-stress preconditioning influences long-term endothelial traction and alignment under high shear flow. American Journal of Physiology-Heart and Circulatory Physiology. 2024;326:H1180–H1192. doi: 10.1152/ajpheart.00067.2024

31. Han SJ, Oak Y, Groisman A, Danuser G. Traction microscopy to identify force modulation in subresolution adhesions. Nature Methods. 2015;12:653–656. doi: 10.1038/nmeth.3430

32. Haarman SE, Kim SY, Isogai T, Dean KM, Han SJ. Particle retracking algorithm capable of quantifying large, local matrix deformation for traction force microscopy. PloS one. 2022;17:e0268614.

33. Mittal N, Han SJ. High-Resolution, Highly-Integrated Traction Force Microscopy Software. Current protocols. 2021;1:e233.

34. Chandurkar MK, Han SJ. Subcellular Force Quantification of Endothelial Cells Using Silicone Pillar Arrays. Methods Mol Biol. 2022;2375:229–245. doi: 10.1007/978-1-0716-1708-3_19

35. Pachitariu M, Stringer C. Cellpose 2.0: how to train your own model. Nature Methods. 2022;19:1634–1641. doi: 10.1038/s41592-022-01663-4

36. Li J, Hou B, Tumova S, Muraki K, Bruns A, Ludlow MJ, Sedo A, Hyman AJ, McKeown L, Young RS, et al. Piezo1 integration of vascular architecture with physiological force. Nature. 2014;515:279–282. doi: 10.1038/nature13701

37. Bhatt A, Kaverina I, Otey C, Huttenlocher A. Regulation of focal complex composition and disassembly by the calcium-dependent protease calpain. Journal of cell science. 2002;115:3415–3425.

38. Sjaastad MD, Lewis RS, Nelson WJ. Mechanisms of integrin-mediated calcium signaling in MDCK cells: regulation of adhesion by IP3-and store-independent calcium influx. Molecular biology of the cell. 1996;7:1025–1041.

39. Hartmannsgruber V, Heyken WT, Kacik M, Kaistha A, Grgic I, Harteneck C, Liedtke W, Hoyer J, Kohler R. Arterial response to shear stress critically depends on endothelial TRPV4 expression. PLoS One. 2007;2:e827. doi: 10.1371/journal.pone.0000827

40. Baratchi S, Knoerzer M, Khoshmanesh K, Mitchell A, McIntyre P. Shear Stress Regulates TRPV4 Channel Clustering and Translocation from Adherens Junctions to the Basal Membrane. Scientific Reports. 2017;7:15942. doi: 10.1038/s41598-017-16276-7

41. Sweet DT, Chen Z, Givens CS, Owens III AP, Rojas M, Tzima E. Endothelial Shc regulates arteriogenesis through dual control of arterial specification and inflammation via the notch and nuclear factor-κ–light-chain-enhancer of activated B-cell pathways. Circulation research. 2013;113:32–39.

42. Gerhold KA, Schwartz MA. Ion channels in endothelial responses to fluid shear stress. Physiology. 2016;31:359–369.

43. Oldham WM, Hamm HE. Heterotrimeric G protein activation by G-protein-coupled receptors. Nature reviews Molecular cell biology. 2008;9:60–71.

44. Webb DJ, Donais K, Whitmore LA, Thomas SM, Turner CE, Parsons JT, Horwitz AF. FAK-Src signalling through paxillin, ERK and MLCK regulates adhesion disassembly. Nat Cell Biol. 2004;6:154–161. doi: 10.1038/ncb1094

45. Tran Q-K, Ohashi K, Watanabe H. Calcium signalling in endothelial cells. Cardiovascular Research. 2000;48:13–22. doi: 10.1016/s0008-6363(00)00172-3

46. Ahn SJ, Fancher IS, Bian J-T, Zhang CX, Schwab S, Gaffin R, Phillips SA, Levitan I. Inwardly rectifying K+ channels are major contributors to flow-induced vasodilatation in resistance arteries. The Journal of Physiology. 2017;595:2339–2364. doi: 10.1113/JP273255

47. Ran W, Stephanie EL, Stephen LG, Paul MK, Frances P. The Endothelium: The Vascular Information Exchange. In: John NB, Erik JB, eds. Calcium and Signal Transduction. Rijeka: IntechOpen; 2018:Ch. 3.

